# NewtCap: an efficient target capture approach to boost genomic studies in Salamandridae (true salamanders and newts)

**DOI:** 10.1101/2024.10.25.620290

**Authors:** Manon Chantal de Visser, James France, Evan McCartney-Melstad, Gary M. Bucciarelli, Anagnostis Theodoropoulos, Howard Bradley Shaffer, Ben Wielstra

## Abstract

Salamanders have large and complex genomes, hampering whole genome sequencing. However, reduced representation sequencing provides a feasible alternative to obtain genome-wide data. We present NewtCap: a sequence capture bait set that targets c.7k coding regions across the genomes of all true salamanders and newts (the family Salamandridae, also known as ‘salamandrids’). We test the efficacy of NewtCap, originally designed for the Eurasian *Triturus* newts, in 30 species, belonging to 17 different genera, that cover all main Salamandridae lineages. We also test NewtCap in two other salamander families. We discover that NewtCap performs well across all Salamandridae lineages (but not in the salamander families Ambystomatidae and Hynobiidae). As expected, the amount of genetic divergence from the genus *Triturus* correlates negatively to capture efficacy and mapping success. However, this does not impede our downstream analyses. We showcase the potential of NewtCap in the contexts of; 1) phylogenomics, by reconstructing the phylogeny of Salamandridae, 2) phylogeography, by sequencing the four closely related species comprising the genus *Taricha*, 3) hybrid zone analysis, by genotyping two *Lissotriton* species and different classes of interspecific hybrids, and 4) conservation genetics, by comparing *Triturus ivanbureschi* samples from several wild populations and one captive-bred population. Overall, NewtCap has the potential to boost straightforward, reproducible, and affordable genomic studies, tackling both fundamental and applied research questions across salamandrids.

## Introduction

One of the most challenging groups of animals to study genomically are the salamanders. These organisms have complex and large genomes that contain many repetitive elements compared to most other animals (e.g. they can be in the range of 10 to 40 times the size of a human genome; Gregory, 2002, Litvinchuk et al., 2007, Sessions, 2008, Sun et al., 2012, Gregory, 2024, Smith et al., 2019). This is the primary reason that conducting whole genome sequencing and *de novo* genome assembly for salamanders is extremely costly in terms of money, time, and computational resources (Lou et al., 2021, Calboli et al., 2011).

Fortunately, reduced representation sequencing strategies are paving the way toward more straightforward, reproducible, and affordable genomic studies, especially in organisms with large and complex genomes (Good, 2011, Zaharias et al., 2020). By focusing sequencing efforts on subsets of the genome, rather than the entire genome, valuable time and resources can be conserved, allowing for a greater number of samples to be processed (Andermann et al., 2019). Different types of genome-subsampling techniques exist, broadly categorized as ‘non-targeted’ versus ‘targeted’, and their suitability varies, depending on the study species and research objectives (Da Fonseca et al., 2016, Lemmon and Lemmon, 2013, Jones and Good, 2016).

Non-targeted approaches, such as Restriction site-Associated DNA sequencing (RAD-seq) and related techniques, are widely used in genetic mapping and population studies, including salamander studies (e.g. Hu et al., 2019, Rancilhac et al., 2019, Hubbs et al., 2022, Andrews et al., 2016, France et al., 2024a, Babik et al., 2024, Rodríguez et al., 2017, Burgon et al., 2021, Scott et al., 2024). While simple, scalable (Andermann et al., 2019), and having the key advantage of not needing to know the sequences of any loci beforehand, such non-targeted approaches are known to yield missing data, underestimate genetic diversity, and call incorrect allele frequencies (Arnold et al., 2013, Rubin et al., 2012, Davey et al., 2013). This is due to restriction site polymorphisms that limit the phylogenetic signal for resolving deep, evolutionary relationships (Dodsworth et al., 2019, Rubin et al., 2012, Lowry et al., 2017).

On the other hand, targeted methods such as target capture sequencing offer a strategy to achieve higher resolution and specificity, as they focus on pre-selected (orthologous) loci (Mamanova et al., 2010, Harvey et al., 2016). With target capture sequencing, biotinylated RNA probes – or ‘baits’ – that are complementary to the loci of interest are used, causing them to bind to the ‘target’ regions, before streptavidin-coated magnetic beads are used to ‘capture’ them (Gnirke et al., 2009, Albert et al., 2007). The main advantage of using the target capture method over untargeted methods is that there will be higher efficiency, because enrichment of specific (usually coding) genomic regions of interest is more effectively achieved across samples (Grover et al., 2012, Andermann et al., 2019, Heyduk et al., 2016, Fitzpatrick et al., 2024). Furthermore, the flanking regions of targets can also provide information on more variable genomic regions such as introns (Jones and Good, 2016, Zhou and Holliday, 2012), and off-target ‘bycatch’ reads can provide additional data as well (Featherstone and McGaughran, 2024, Guo et al., 2012).

Often custom target capture baits are designed, which can be based on a draft genome or transcriptome reference, however pre-designed baits can be ordered as well (Bi et al., 2012, Andermann et al., 2019, Jimenez-Mena et al., 2022). Over the last decade, many target capture bait sets have been tested and made publicly available for different types of organisms, ranging from micro-organisms to macro-organisms (e.g.; Andermann et al., 2019, Grover et al., 2012, Heyduk et al., 2016, Khan et al., 2024, Quek and Ng, 2024, Yu et al., 2023). For animals in particular, bait sets exist for certain groups of insects (e.g. Hymenoptera; Faircloth et al., 2015), snails (e.g. Eupulmonata; Teasdale et al., 2016), reptiles (e.g. Squamata; Schott et al., 2017, Singhal et al., 2017), fish (e.g. Acanthomorpha; Alfaro et al., 2018), and amphibians (e.g. Anura; Hutter et al., 2022). Salamander bait sets have also already been designed for particular genera (*Ambystoma* and *Triturus*; McCartney-Melstad et al., 2016, Wielstra et al., 2019). As the target capture approach generally handles a certain degree of sequence divergence well, a bait set designed for one genus has great potential to work in other genera too (Bragg et al., 2016, Portik et al., 2016). Thus, it is worth exploring the potential transferability of such existing bait sets to related taxa.

We introduce ‘NewtCap’: a target capture bait set and protocol for salamanders of the family Salamandridae (which includes the ‘true salamanders’ and the ‘newts’). The bait set, originally designed for *Triturus* newts (Wielstra et al., 2019), targets c. 7k putative, orthologous, coding regions. Here, we provide an updated version of the lab protocol that cuts down on costs, time and DNA input. Furthermore, we investigate the efficacy of NewtCap across 30 different species and 17 genera of the Salamandridae family and test its efficiency in two distantly related species of other salamander families. Besides checking general performance, we assess the usefulness of NewtCap in the contexts of; 1) phylogenomics, by inferring a Salamandridae phylogeny, 2) phylogeography, by investigating the relationships among multiple individuals of four closely related *Taricha* species, 3) hybrid zone analysis, by calculating the hybrid index and heterozygosity for two *Lissotriton* species and interspecific hybrids of different cross types, and 4) conservation genetics, by determining the genetic relatedness of a captive-bred *Triturus ivanbureschi* population to wild populations. Overall, we demonstrate that NewtCap is an important tool for molecular studies on salamandrids.

## Materials and Methods

### Sampling and DNA Extraction

We studied 73 individual salamanders in total, covering 30 species and 17 genera of the Salamandridae family (including nine interspecific hybrids of the genus *Lissotriton*), as well as two more distantly related samples (Table S1, see <u>Zenodo</u>) from the families Ambystomatidae and Hynobiidae (Marjanović and Laurin, 2013, Frost, 1985). We obtained DNA extractions or genetic data from previous studies (Wielstra et al., 2019, De Visser et al., 2024, Kazilas et al., 2024, Mars et al., 2025, Kalaentzis et al., 2025, Robbemont et al., 2023, Pasmans et al., 2006, Valbuena-Urena et al., 2013, Sequeira et al., 2022, France et al., 2024b), as well as new samples for this study (provided by collaborators, see acknowledgments and Table S1 via <u>Zenodo</u>).

We used the Promega Wizard™ Genomic DNA Purification Kit (Promega, Madison, WI, USA), which is a salt-based DNA extraction protocol (Sambrook and Russell, 2001). The DNA from each sample was re-suspended in 100 µL 1 × TE buffer before concentrations and purity were assessed via spectrophotometry, using the DropSense96™ (Trinean, Gentbrugge, Belgium). As we used a minimum concentration of 150 ng/µL for library preparation, any sample found to be below was concentrated by vacuum centrifugation.

#### NewtCap: Laboratory procedures

The target capture procedures are fully documented and included as Supplementary Materials (see under ‘Data Accessibility’). These protocols are rigorously optimized versions of those previously described (see Wielstra et al., 2019). Sonication of genomic DNA has been replaced with enzymatic fragmentation, resulting in a tenfold increase in library yield per ng of input DNA, in addition to time and cost savings. The volumes of all reagents in the library preparation were reduced by 75% compared to the manufacturers protocol to conserve reagents. Target capture protocols have been adapted from Mybaits V3.0 to the V4.0 kit and are fully compatible with Mybaits V5.0.

### Library preparation

DNA libraries were constructed using the NEBNext Ultra™ II FS DNA Library Prep Kit for Illumina (New England Biolabs, Ipswich, MA, USA) following the manufacturers protocol, with quarter volumes of all reagents and with the following modifications: 1,000 ng of extracted genomic DNA was used as input (6.5 µL at 154 ng/µL). The enzymatic shearing time at 37 °C was adjusted to 6.5 minutes (as the minimum time of 15 minutes suggested by the manufacturer resulted in over-digestion). After NEB adapter ligation and cleavage with the NEB USER enzyme, NucleoMag™ magnetic separation beads (Macherey-Nagel, Düren, Germany) were used for double-ended size selection targeting an insert size of 300 bp. Libraries were indexed with 8 cycles of PCR amplification, using unique combinations of custom i5 and i7 index primers (Integrated DNA Technologies, Leuven, Belgium). NucleoMag™ beads were used again for a final cleanup before the libraries were resuspended in 22 µL of 0.1 × TE buffer. Library size distribution and concentration was measured using the Agilent 4150 TapeStation or 5200 Fragment analyzer system (Agilent Technologies, Santa Clara, CA, USA), using the D5000 ScreenTape or DNF-910 dsDNA Reagent Kit. We aimed for obtaining a final library concentration of at least 12 ng/µL.

### Target Capture, Enrichment, and Sequencing

Libraries were equimolarly pooled in batches of 16, aiming for a total DNA mass of 4,000 ng (250 ng per sample). Vacuum centrifugation was then used to reduce the volume of each pool to 7.2 µL (556 ng/µL). We performed target capture with the MyBaits v4.0 kit (Arbor Biosciences, Ann Arbor, MI, USA) previously designed for *Triturus* newts (Wielstra et al., 2019), which targets 7,139 unique exonic regions (product Ref: # 170210-32). The manufacturers protocol was employed with the following deviations: Blocks C and O were replaced with 5 µL of *Triturus* derived C0t-1 DNA at 6,000 ng/µL (30,000 ng per pool). C0t-1 DNA is enriched in repetitive sequences and acts as a blocking buffer to non-specific targets in capture assays by hybridizing with repetitive sequences in the libraries (McCartney-Melstad et al., 2016). Tissue to produce C0t-1 DNA was derived from an invasive population of *T. carnifex* (Meilink et al., 2015).

The pooled libraries were incubated with the blocking buffer for 30 minutes, followed by hybridization for 30 hours at 62 °C. After capture of the hybridized baits with streptavidin coated beads and four cycles of washing, each pool was divided into equal-volume halves. The first half was subject to 14 cycles of PCR amplification, followed by bead cleanup, and resuspension in 22 µL of 0.1 × TE buffer. The concentration and fragment size distribution of the enriched pool were then measured with the TapeStation system, using the HS D5000 ScreenTape kit. If the final concentration was between 5 and 20 nM then the same protocol was employed for the second half of the pool. If not, the number of post-enrichment PCR cycles was altered to compensate. For each pool 16 GB (1 GB per sample) of 150 bp paired-end sequencing was performed on the Illumina NovaSeq 6000 platform (Illumina Inc., San Diego, CA, USA) by BaseClear B.V. (Leiden, the Netherlands).

#### Bioinformatics for data pre-processing

We followed a standard pipeline for cleaning up the raw reads and mapping the trimmed sequence data against reference sequences in a Linux environment (adapted from Wielstra et al., 2019). We describe the main steps and provide the scripts we used publicly through GitHub: https://github.com/Wielstra-Lab/NewtCap_bioinformatics.

### Quality Control and Read Clean-up

First, we clipped all reads to a maximum length of 150 bp using the BBDuk script by BBTools (BBMap – Bushnell B. –). Then we removed any leftover adapter contamination and low-quality bases (or reads) using Trimmomatic v.0.38 (Bolger et al., 2014). Adapter sequences for the TruSeq2 multiplexed libraries were identified and removed. Leading and trailing bases were trimmed if the Phred score was < 5. We also removed reads in case the average Phred score in a sliding window (5’ to 3’) of a size of five base pairs dropped below 20, and we discarded all reads shorter than 50bp. We monitored the quality of our sequences before and after trimming with FastQC (Andrews, 2010) by using the summarizing ‘quality_check’ function of SECAPR (Andermann et al., 2018).

### Read Mapping and Variant Calling

Cleaned reads were mapped to the set of 7,139 *T. dobrogicus* reference sequences with a maximum length of 450bp (based on *T. dobrogicus* transcripts, see Wielstra et al., 2019) that were initially used for probe development. The reference FASTA file is provided as Supplementary Material. Mapping was performed using the MEM algorithm from Burrows-Wheeler Aligner v.0.7.17 (Li, 2013), and we stored results in BAM format using SAMtools v.1.18 (Li et al., 2009, Danecek et al., 2021). We added read group information using the AddOrReplaceReadGroups function of Picard v.2.25.1 and PCR duplicates were flagged with Picard’s MarkDuplicates (http://broadinstitute.github.io/picard/). After doing so, the output was again saved in BAM format.

Next, we called variants using the HaplotypeCaller function of GATK v.4.1.3.0 (Auwera and O’Connor, 2020) including the -ERC GVCF option, and we used GATK’s CombineGVCFs and GenotypeGVCFs functions to perform joint genotype calling to create multi-sample (ms) gVCF files as input for downstream analyses. In case samples still needed to be added or removed from msgVCF files after joint genotyping, we did so using the view function of BCFtools v.1.15.1 (Danecek et al., 2021, Li, 2011).

#### Downstream Analyses

### Bait performance statistics

To evaluate the overall performance of the NewtCap baits, we calculated several statistics (Table S1). First, we counted the number of GB and reads contained in the raw FASTQ files for each sample. We also used the SAMtools’ flagstat function to determine the total number of reads present in the (deduplicated) BAM files that passed quality control, as well as the percentage of these reads that successfully mapped to a reference sequence. We used SAMtools’ coverage function to extract information on the mean depth of coverage, as well as the mean percentage of coverage, for each target, and then averaged these per sample to provide an estimate of the “success rate”. Besides analyzing the contents of the BAM files, we checked how many sites were outputted in total in each separate, raw gVCF file by counting lines (as each non-header line is one site). Then, we counted how many of those were considered SNPs, and how many were considered INDELs, by using the stats function of BCFtools (note that this is done before merging any files or applying any SNP filtering).

To estimate how the performance of NewtCap correlates with the level of genetic divergence from *T. dobrogicus*, the species that was used for bait design and as a reference for read mapping, we performed several statistical analyses. For these, we used a rough estimate of divergence times for lineages of Salamandridae (from De Visser et al., 2024). We explored the relationship between this estimated genetic divergence from *T. dobrogicus* and the following performance variables; 1) the percentage of mapped reads, 2) the percentage of reads marked as PCR duplicates, 3) the mean read depth number after deduplication, 4) the mean coverage of the sequence bases after deduplication, 5) the number of SNPs found in the raw gVCFs files, and 6) the number of INDELs found in the gVCF files. We calculated the correlation coefficients (r) between either of these variables and the estimated genetic divergence from *T. dobrogicus*. Depending on whether our data met the assumptions for parametric or non-parametric testing, we either employed the Pearson correlation method or the Spearman’s rank correlation method. We determined the level of significance using a two-tailed test, with the p-value threshold of p<0.00833 to indicate statistical significance (implementing a Bonferroni correction on the usual threshold p<0.05 for the six tests performed, as 0.05/8=0.00833). These analyses, including assumption checks such as testing for normality, were performed in Microsoft Excel 2024 (https://office.microsoft.com/excel).

### Concatenated phylogenetic analyses in RAxML

To investigate the usefulness of NewtCap in the context of phylogenomics, we built phylogenetic trees with a Maximum Likelihood method using RAxML. We used NewtCap to reconstruct an existing Salamandridae tree that was originally built based on 5,455 nuclear genes derived from transcriptome data (Rancilhac et al., 2021), by including 23 samples that cover at least one representative genus for each of the main clades *in sensu* (as described in Rancilhac et al., 2021). To further check the performance of NewtCap across true salamanders as well as for non-salamandrid species, and to explore the position of the root, we added three additional samples in a second, extended analysis, namely the Salamandridae species *Mertensiella caucasica* (family Salamandridae) and two non-salamandrid species *Ambystoma mexicanum* (family Ambystomatidae) and *Paradactylodon gorganensis* (family Hynobiidae). The raw msgVCF files for these two analyses respectively comprised 812,574 sites and 812,603 sites from across 7,135 targets. Note that not all sites in the raw and intermediate gVCF and msgVCF files were necessarily polymorphic (i.e. invariant - or monomorphic - sites may still have been included, unless we specifically stated that these have been removed).

We applied quality filtering on the msgVCFs (which contained only the samples chosen for these phylogenetic analyses). First, we removed sites that showed heterozygote excess (p<0.05) using BCFtools v.1.15.1, in order to rid paralogous targets (see Wielstra et al., 2019). Then, we used VCFtools v.0.1.16 to enforce the following strict filtering options; discarding INDELs (“--remove-indels”), filtering out sites with for instance poor genotype and mapping quality scores (“QD<2”, “MQ<40”, “FS>60”, “MQRankSum <−12.5”, “ReadPosRankSum<−8”, and “QUAL<30”; Poplin et al., 2017), and discarding sites with over 50% missing data (“--max-missing=0.5”). At this stage, the intermediate msgVCF files contained, in total, 625,182 SNPs from across 7,103 targets, and 621,754 SNPs from across 7,095 targets, for the sets of 23 and 26 samples.

We converted the files into PHYLIP format with the ‘vcf2phylip.py’ script (Ortiz, 2019), which, by default, requires a minimum of four samples representing each SNP variant. Next, we performed an ascertainment bias correction by using the ‘ascbias.py’ script (https://github.com/btmartin721/raxml_ascbias) to remove sites considered invariable by RAxML. The final PHYLIP file was used as input for RAxML v.8.2.12 (Stamatakis, 2014), which we ran with 100 rapid bootstrap replicates, under the ASC_GTRGAMMA model, and with the Lewis ascertainment correction (Lewis, 2001). We obtained the best-scoring Maximum Likelihood tree out of the concatenation analyses.

To gain insights into the potential of NewtCap in a phylogeographical context, we also built a phylogenetic tree for the New World newt genus *Taricha* using RAxML. As input, we used a msgVCF file containing nine samples: eight *Taricha* samples covering four distinct species within the genus (*T. granulosa*, *T. rivularis*, *T. sierrae* and *T. torosa*, with two samples per species) and one sample of the sister-genus *Notophthalmus* as outgroup (Table S1). The raw msgVCF file comprised 209,072 sites from 7,059 targets. Subsequently, we performed the filtering steps as described above: rusing BCFtools v.1.14.1 to remove heterozygote excess (p<0.05), and using VCFtools v.0.1.16 to discard INDELs (“--remove-indels”), filter out sites with poor quality scores (“QD<2”, “MQ<40”, “FS>60”, “MQRankSum <−12.5”, “ReadPosRankSum<−8”, and “QUAL<30”), and discard sites with over 50% missing data (“--max-missing = 0.5”). This left the intermediate msgVCF file with 180,385 SNPs from across 7,048 targets. We visualized all phylogenies using FigTree v.1.4.4 (http://tree.bio.ed.ac.uk/software/figtree/).

### Hybrid analyses and conservation genetics

To assess the usefulness of NewtCap in the context of hybridization studies, we used the R packages ‘*triangulaR’* (Wiens and Colella, 2024), ‘*ggplot2’* (Wickham, 2011) and ‘*vcfR’* (Knaus and Grunwald, 2017) to build triangle plots. We investigated the target capture data of in total 15 *Lissotriton* individuals; three of each parental species (*L. vulgaris* and *L. montandoni*), as well as hybrids bred and reared in the lab: three F1 hybrids, three F2 hybrids, and three backcrosses (F1 x *L. vulgaris*). We assembled a raw msgVCF file containing only these samples, which initially had data on 807,389 sites across in total 7,135 targets. From this we extracted high quality SNPs as described before (by removing heterozygote excess, discarding indels, and filtering out sites with poor quality scores), except here we tolerated no missing data at all (i.e., we set VCFtools’ “--max-missing” option to 1.0, instead of 0.5). The filtered msgVCF contained 532,332 SNPs from 6,866 targets in total and was used as input for *triangulaR*.

The *triangulaR* package uses SNPs from an input VCF file that are estimated to be species-diagnostic based on the parent species under a certain allele frequency threshold between ‘0’ and ‘1’, where ‘1’ equals a fixed difference between parental population (Wiens and Colella, 2024). Thus, the threshold of 1 is employed when searching for two consistent, diverged, homozygous states in the parental populations. SNPs that pass this filter can then be used to calculate the hybrid index and heterozygosity for the samples analyzed. Considering that our parent populations did not comprise the actual parents of the hybrids included in our study and were represented by just three samples each, this complicated distinguishing species-diagnostic SNPs from SNPs that only appeared diagnostic by chance. Therefore, we only extracted SNPs that were 100% species-diagnostic (based on the information from the parental populations, by setting the allele frequency threshold to 1) *and* that were always heterozygous in the F1’s. This extra functionality (i.e. filtering for heterozygosity in F1 hybrids) is currently not built into *triangulaR*, but we added this filtering option before conducting our calculations. We provide the customized R script in our GitHub repository.

To assess the performance of NewtCap in the context of a conservation genetic study we conducted a Principal Component Analysis (PCA) and Hierarchical Clustering Analysis (HCA) to determine the geographical origin of a captive bred population, which is presumed to originate from Cerkezköy in Turkey (Michael Fahrbach, pers. comm.). We used the R packages ‘*gdsfmt’* and ‘*SNPRelate’* (Zheng et al., 2012) on data from 24 *Triturus ivanbureschi* newts that originated from seven different wild populations and the captive population (i.e. three samples per population, see Table S1). Populations from the wild include both the glacial refugial area (Asia Minor and Turkish Thrace) and postglacially colonized area (the Balkans; Wielstra et al., 2017). We created a raw msgVCF file from these samples, which contained a total of 812,768 sites from across 7,135 targets. We extracted high quality SNPs in the same way as we did for the hybrid studies (i.e. removing heterozygote excess, discarding indels, filtering out sites with poor quality scores, and allowing no missing data), and the filtered msgVCF that was used as input had a total of 486,891 SNPs from 5,774 targets. We conducted the hybrid and conservation genetic analyses in Rstudio (Team_R_Studio, 2021) using R v.4.1.2 (Team_R_Core, 2020).

Finally, we were interested in the number of informative SNPs that we could find within Salamandridae species that are distantly related to *T. dobrogicus.* The most distantly related species for which we had more than one sample belonging to distinct populations in our dataset is *Chioglossa lusitanica* (Table S1), which is a true salamander, not a newt. Having more than one sample allows us to cross-compare these populations. Thus, we extracted the variant information of the two *C. lusitanica* samples from the overall msgVCF file –filtered for high quality SNPs allowing no missing genotypes– that was used to estimate the genetic divergence from *T. dobrogicus* (De Visser et al., 2024). We quantified the number of SNPs that were homozygous in one sample and heterozygous in the other sample – and *vice versa*. Additionally, we counted the number of SNPs that were homozygous in both samples, but for different alleles.

## Results

### Overall performance of NewtCap

In total we collected 149.0 GB of raw sequence data with on average 15,573,781 reads per sample (s.d.=11,400,957, see Table S1 for the read counts per sample). On average 42.6% (s.d.=16.2%) of all reads were successfully mapped against the *Triturus* reference, and an average of 11.9% (s.d.=10.6%) of all reads were marked as PCR duplicates. In the deduplicated BAM files, the mean read depth across all targets per sample varied widely, and ranged from 5.9 X (s.d.=27.9) in *A. mexicanum* and 8.2 X (s.d.=27.8) in *P. gorganensis* to 122.2 X (s.d.=345) in one of the *Lissotriton* hybrids. The mean coverage of the sequence bases of all targets ranged from 36.1% (s.d.=30.3%) in *P. gorganensis* and 36.4% (s.d.=31.3%) in *A. mexicanum* to 98.31% in two *T. ivanbureschi* samples (s.d.=7.2% and s.d.=6.9%). On average, the mean read depth was 60.5 X (s.d.=183.2) across all samples, and the overall mean coverage of the sequence bases was 92.0% (s.d.=13.7%). Details can be found in Table S1.

We discovered significant correlations between the genetic divergence from *T. dobrogicus* and several performance variables. Firstly, we found no significant correlation between the percentage of reads marked as PCR duplicates by Picard and the level of divergence from *T. dobrogicus* (Spearmann’s rank correlation test, R=0.015, p=0.91), however we did find a negative correlation between the percentage of reads that map against the *T. dobrogicus* reference sequences and the amount of divergence from *T. dobrogicus* (Spearmann’s rank correlation test, R=-0.661, p<0.001, Table 1). Furthermore, there is a negative correlation between the coverage of reference targets and the level of divergence from *T. dobrogicus* - i.e. the mean read depth and the mean coverage of the sequence bases both drop as the divergence from *T. dobrogicus* increases (Pearson correlation test for the mean read depth, R=-0.597, p<0.001 and Spearmann’s rank correlation test for the mean coverage of the sequence bases, R=-0.918, p<0.001; Table 1). Lastly, with increased divergence from *T. dobrogicus*, more SNPs and INDELs are discovered by BCFtools in the eventual gVCF files (Spearmann’s rank correlation test; R=0.917 and p<0.001 for SNPs, R=0.899 and p<0.001 for INDELs, Table 1).

**Table 1:** Correlation statistics for the relationship between NewtCap performance variables and the estimated amount of species divergence from Triturus dobrogicus. For each of the separate performance variable the appropriate statistical test - either the Spearmann’s rank or the Pearson correlation test - was applied to determine whether the observed correlations were significant. Both the correlation coefficient (R) and its square value are provided, as well as the p-value. Results with a p-value lower than 0.008335 (including a Bonferroni correction, see Methods) are marked with an asterisk.

| Performance Variable | Test applied | R | R<sup>2</sup> | P-value |
| --- | --- | --- | --- | --- |
| Percentage of reads mapped | Spearmann | -0.661 | 0.440 | 3.6E-10* |
| Percentage of PCR duplicates | Spearmann | 0.015 | < 0.001 | 0.910 |
| Mean read depth | Pearson | -0.597 | 0.357 | 3.8E-08* |
| Mean coverage of sequence bases | Spearmann | -0.918 | 0.842 | 2.2E-29* |
| Number of SNPs discovered | Spearmann | 0.917 | 0.841 | 2.7E-29* |
| Number of INDELs discovered | Spearmann | 0.899 | 0.807 | 2.4E-26* |

### Phylogenomics: Reconstruction of the Salamandridae phylogeny

The fully resolved and highly supported Salamandridae phylogeny resulting from Maximum Likelihood analysis of concatenated data in RAxML was based on 204,600 polymorphic SNPs (Fig. 1 and Fig. S1). We recover the newts and the true salamanders as monophyletic groups. Within the newts, the modern European and modern Asian newts are sister lineages. Their closest relatives are the Corsica/Sardina newts. The New World newts are the sister lineage of this assemblage. Finally, these newt lineages coalesce with the primitive newts at the newt crown. The placement of the root on the branch connecting the newts and the clade containing the true salamanders and *Salamandrina* was confirmed by our extended phylogeny (based on 265,105 SNPs, Fig. S2) that included two non-Salamandridae salamander species (*A. mexicanum* and *P. gorganensis*), as well as an additional true salamander species (*M. caucasica*, which is recovered as the sister lineage of *Chioglossa*).

**Figure 1:**
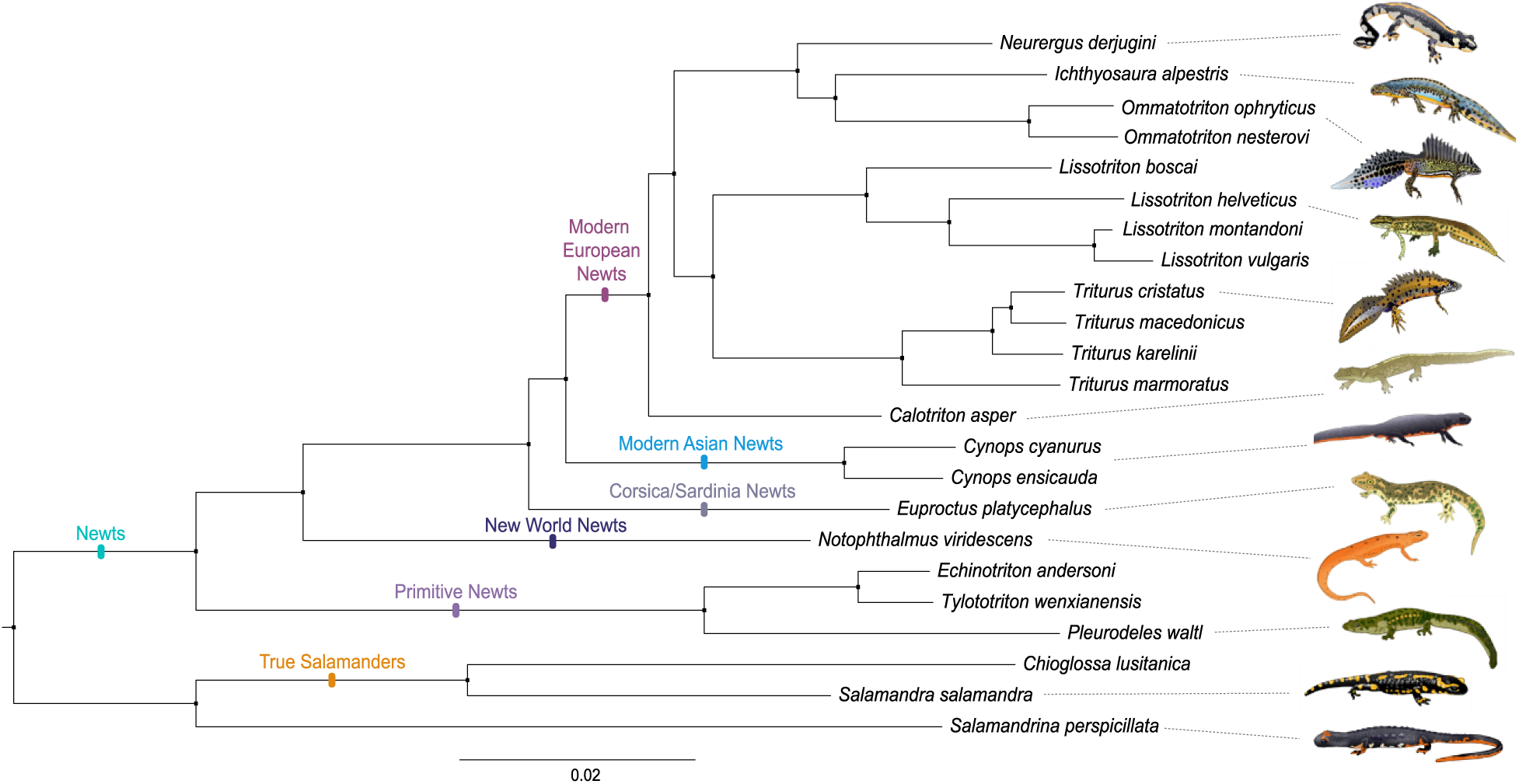
NewtCap-based phylogeny of the Salamandridae family. The phylogeny is based on Maximum Likelihood inference of concatenated data of 204,600 informative SNPs using RAxML. Overall layout and clade labels conform to a previous transcriptome-based phylogeny (Rancilhac et al., 2021). The tree is rooted on the branch separating the newts and the clade containing the true salamanders and Salamandrina (see also Fig. S1 for the same tree, but with original labels, and Fig. S2 for the additional, extended tree, including Ambystoma, Paradactylodon and Mertensiella, that confirms the root position adopted here). All nodes have a bootstrap support of 100%.

### Phylogeography: Fully resolved relationships of *Taricha*

The *Taricha* phylogeny that resulted from the concatenated data used by RAxML to perform a Maximum Likelihood analysis, was based on 9,730 polymorphic SNPs. The tree was fully resolved and highly supported (Fig. 2). All four *Taricha* species are recovered as reciprocally monophyletic. The basal bifurcation is between *T. granulosa* and the remainder. The subsequent split separates *T. rivularis* from *T. torosa* plus *T. sierrae*.

**Figure 2:**
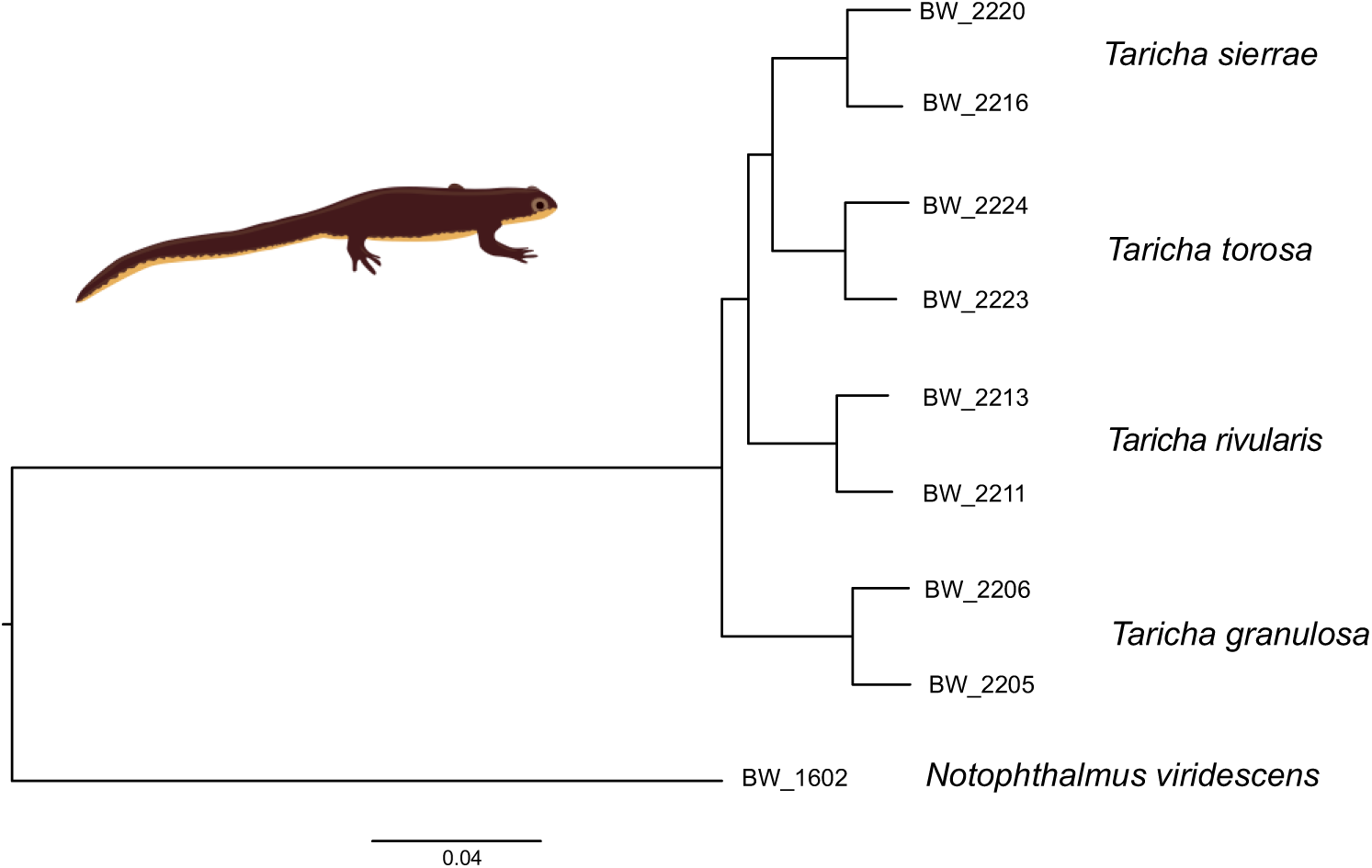
A Taricha phylogeny obtained with NewtCap-derived data. The phylogeny is based on Maximum Likelihood inference of concatenated data of 9,730 informative SNPs using RAxML. Notophthalmus *is used to root the tree. All nodes have a bootstrap support of 100% (not shown)*.

### Hybridization studies: *Lissotriton* hybrids and backcrosses detected

After quality filtering, we identified 666 SNPs that we consider species-diagnostic (i.e. ‘ancestry-informative’) for *L. vulgaris* versus *L. montandoni* in the target capture data based on the genotypes of the six parental species samples and three F1 hybrids. Those SNPs enabled us to calculate the hybrid indices and interclass heterozygosity values in F2 and backcross (‘Bx’) hybrids, as visualized in a triangle plot (Fig. 3). The backcross hybrids are placed along the edge of the triangle plot towards the appropriate parental species (*L. vulgaris*) whereas the F2 hybrids are placed more towards the center of the triangle plot.

**Figure 3:**
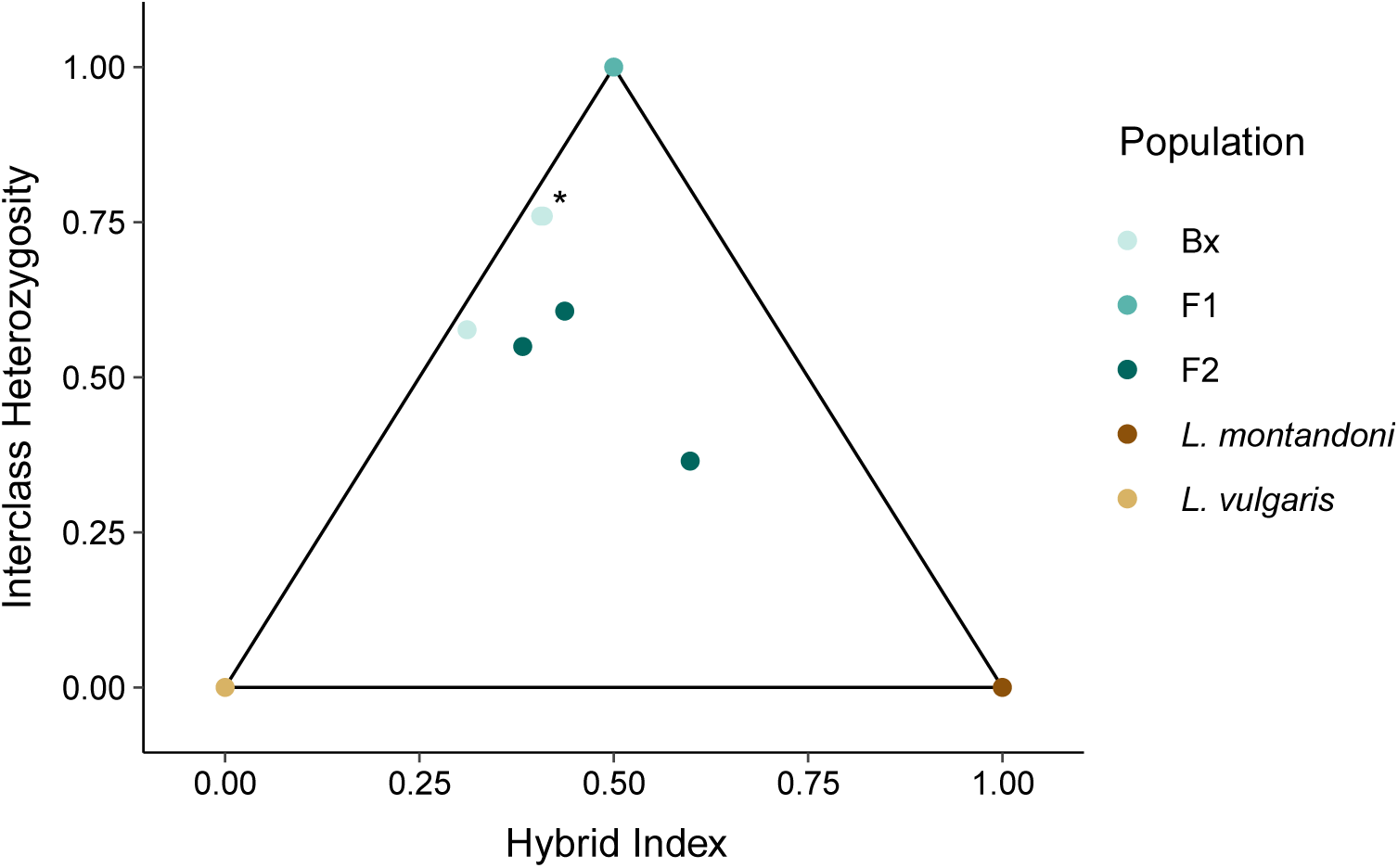
Triangle plot of different Lissotriton hybrid classes based on NewtCap-derived data. The plot, based on 666 informative SNPs, shows the relationship between the hybrid index (the fraction of the alleles per individual that derived from each of the two parental species, also known as the ancestry) and the interclass heterozygosity (the fraction of the alleles per individual that is heterozygous for alleles from both parental species). The L. vulgaris *individuals are in the bottom left corner, the* L. montandoni *individuals in the bottom right corner, and the F1 hybrid offspring in the top corner. The F2 and Bx (‘backcross’) hybrids are placed inside the triangle, with two Bx samples almost fully overlapping (marked with *)*.

### Conservation genetics: Separation of *Triturus* populations and *Chioglossa* subspecies

For the PCA and HCA analyses, the R calculations ended up being based on 9,135 bi-allelic SNPs. Along the first and second Principal Components of the PCA, the four wild populations from the postglacially colonized area cluster together, whereas the three wild populations from the glacial refugial area stand relatively apart Fig. 4A & B). The captive-bred individuals cluster closest to the Safaalan group, a population in Turkey just west of Istanbul, close to the presumed source locality Cerkezköy. The same pattern is observed in the HCA dendogram (Fig. 4C).

**Figure 4:**
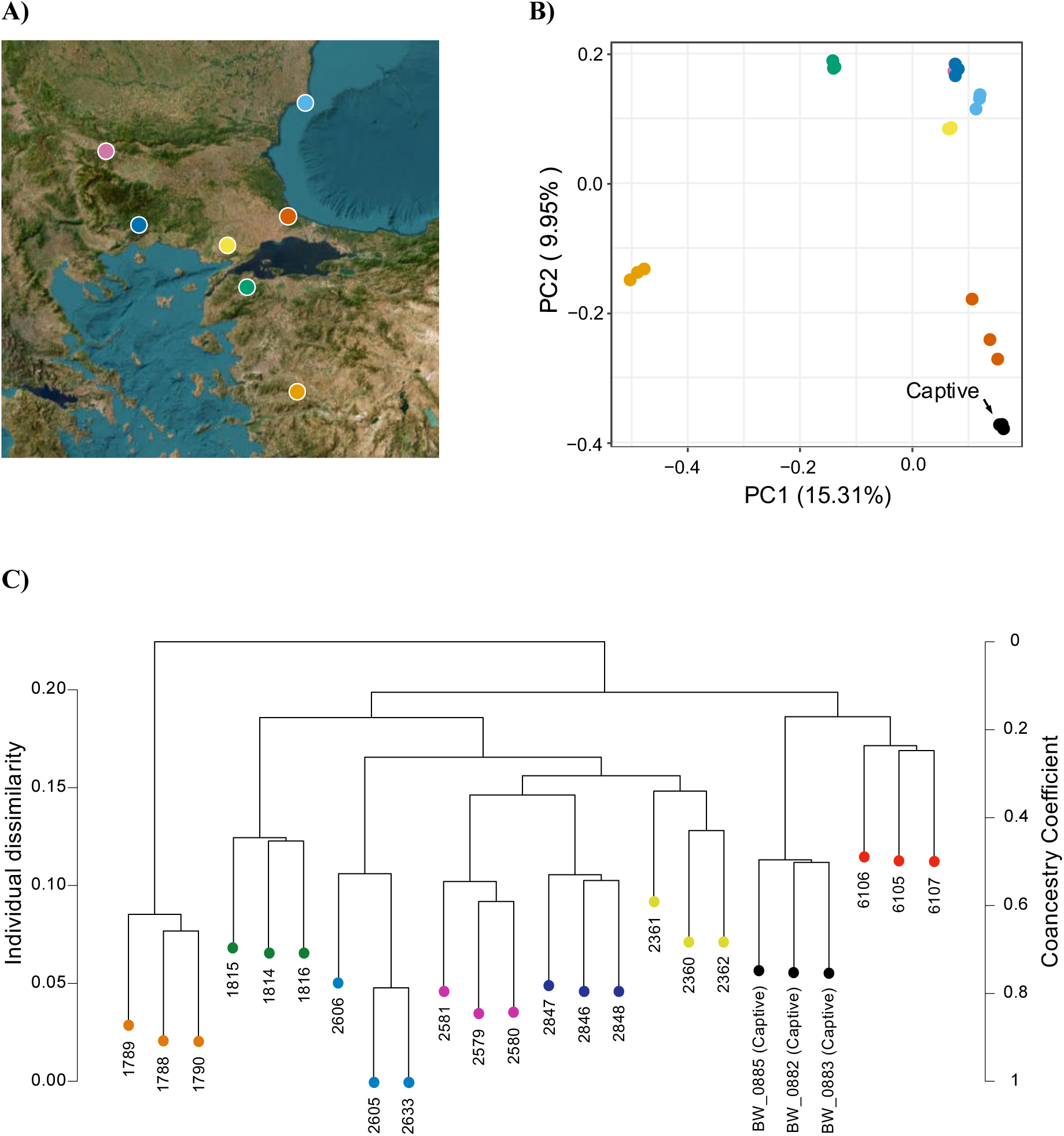
Genetic differentiation between wild and captive Triturus ivanbureschi populations based on NewtCap-derived data. **(A)** The wild population localities (details in Table S1; four postglacial populations are represented by dark blue, light blue, yellow, and pink colors in Bulgaria, Greece and Turkey, and three populations from the glacial refugial area are represented by red, orange and green colors in Turkey). **(B)** A plot of the first versus the second Principal Component (PC) places the captive individuals closest to a population from just west of Istanbul. **(C)** The dendrogram produced by the HCA analysis, showing the Individual Dissimilarity as well as the Coancestry Coefficient, again shows that captive samples cluster with a population just west of Istanbul.

The check for high quality SNPs that display different genotypes in the two *C. lusitanica* samples originating from different subspecies, resulted in a list of over 10,000 polymorphic SNPs: we counted 8,029 SNPs for which one individual was homozygous, and the other individual was heterozygous, and we discovered an additional 2,301 informative SNPs where the individuals were both homozygous, but for alternate alleles.

## Discussion

We introduce NewtCap, a target capture bait and reference set of 7,139 sequences applicable to salamandrids. We show that NewtCap works effectively across all main lineages within the Salamandridae family. As anticipated, the target capture performance and mapping successes are influenced by the level of genetic divergence from *T. dobrogicus* and we do see that – within Salamandridae – more off-target regions are getting captured for more distantly related species. However, these influences appear to be minor and evidently do not hamper our downstream analyses: only for salamanders outside of the family Salamandridae does NewtCap provide data of insufficient quality. NewtCap thus proves to be a powerful tool to collect genomic data for Salamandridae studies regarding systematics (e.g. phylogenomics, phylogeography) and population genetics (e.g. hybrid studies, conservation genetics).

We encourage potential users to further tweak the NewtCap workflow. For instance, in terms of the laboratory protocol, researchers could consider; 1) replacing the C0t-1 blocker with a standard blocker, 2) reducing the hybridization time, and 3) including more individuals per capture reaction. In terms of bioinformatics, the lower mapping rate observed in Salamandridae species more distantly related to *T. dobrogicus* does not necessarily only reflect decreased enrichment efficiency due to genetic divergence; mapping rate is presumably also influenced by the reduced ability of the read mapper to align sequencing reads to more divergent reference sequences used in the bioinformatics pipeline (Bragg et al., 2016, Andermann et al., 2018). Users could explore applying different mapping settings, using alternative mapper tools, or making use of (or generating) substitute reference sequences for read alignment (Schilbert et al., 2020, Andermann et al., 2019).

Our findings demonstrate that the current NewtCap protocol can effectively be applied to any member of the family Salamandridae – the crown of which is dated as far back as c. 100 MYA (Steinfartz et al., 2007, De Visser et al., 2024). Our Salamandridae phylogeny perfectly matches the topology of a transcriptome-based phylogeny (Rancilhac et al., 2021). Although a higher number of molecular markers does not necessarily result in a more accurate species tree (Mirarab et al., 2024, Zhang et al., 2021), independent RAxML analyses using different subsets of NewtCap data result in the same topology (with one notable exception, see; De Visser et al., 2024, France et al., 2024b).

NewtCap has already recently been applied to study the systematics and taxonomy of certain modern European newts – including the genera; *Triturus* (Wielstra et al., 2019, Kazilas et al., 2024), *Lissotriton*, (Mars et al., 2025), *Neurergus* (Koster et al., 2025) and *Ommatotriton*, (Kalaentzis et al., 2025). However, we show that NewtCap allows for a (putative) genome-wide subsampling of thousands of markers for other Salamandridae lineages as well. Existing phylogenies of modern Asian newts (genera *Cynops*, *Paramesotriton*, and *Pachytriton*; Fig. 1) rely on a limited amount of molecular markers and frequently fail to recover genera within the clade as monophyletic groups, presumably due to the intricate biogeographical history of this clade (Veith et al., 2018, Zhang et al., 2008, Kieren et al., 2018). The New World newts (genera *Taricha* and *Notophthalmus*; Fig. 1) have so far only been studied based on mtDNA and allozyme data (Lawson and Kilpatrick, 2014, Gabor and Nice, 2004, Whitmore et al., 2013, Kuchta and Tan, 2006, Kuchta and Tan, 2005). Our *Taricha* phylogeny, the first one based on a considerable number of nuclear markers/SNPs, comprises a fully supported positioning of each of the four known species, with a topology that matches that of an existing mtDNA-based phylogeny (Tan and Wake, 1995, Tan, 1993). For the Asian members of the primitive newts (genera *Tylototriton* and *Echinotriton*; see Fig. 1), genetic resources have been scarce so far, which is why there is an outstanding call to conduct more extensive, genomic research in order to better understand the evolution and taxonomy of the (sub)species of these lineages (Dufresnes and Hernandez, 2022, Weisrock et al., 2006).

NewtCap performs well even for the sister clade of the newts, containing the ‘true salamanders’ (*Salamandra, Chioglossa, Mertensiella, Lyciasalamandra*; see Fig. 1) and the spectacled salamanders (*Salamandrina*). First, NewtCap provides empirical support for the recent suggestion that *Salamandrina* represents the sister lineage to the true salamanders instead of to all remaining salamandrids (Rancilhac et al., 2021). *Salamandrina* itself has so far only been studied with a relatively small number of markers (Mattoccia et al., 2011, Mattoccia et al., 2005, Canestrelli et al., 2014, Hauswaldt et al., 2014, Veith et al., 2009, Weisrock et al., 2006, Zhang et al., 2008). *Chioglossa* is a monotypic genus that is close to becoming threatened according to the IUCN Red List (IUCN_SSC_Amphibian_Specialist_Group, 2022). Conservationists generally identify monotypic taxa as ‘evolutionarily unique’, which helps justify elevated conservation imperatives (Liu et al., 2021, Vane-Wright et al., 1991, Vitt et al., 2023). We obtained c. ten thousand informative SNPs distinguishing two individuals belonging to different *C. lusitanica* subspecies (Sequeira et al., 2022). The sister genus of *Chioglossa*, *Mertensiella*, is monotypic as well, but so far few populations have been studied and only with mtDNA (Tarkhnishvili et al., 2000, Weisrock et al., 2001).

Next to accentuating the potential of NewtCap in the context of conservation genetics, we emphasize that the tool can also be used to identify the degree of interspecific gene flow and geographical structuring. For example, to test hypotheses about species status and historical biogeography. Introgressive hybridization is increasingly recognized as a source of adaptive variation in natural populations, (Fijarczyk et al., 2018, Li et al., 2016, Mallet, 2005), but also as a source of genetic pollution in the case where non-native and native individuals interbreed in nature (Quilodran et al., 2020, Wielstra et al., 2016, Simberloff, 2013, Dufresnes et al., 2015). We identify different hybrid classes of *Lissotriton* newts and are able to differentiate between genetically distinct groups of *T. ivanbureschi* – and determined to which wild population a known, captive population is genetically most similar. Such applications are valuable for guiding wildlife management practices and conservation efforts that concern Salamandridae species both *in situ* and *ex situ* (Frankham et al., 2004, Ørsted et al., 2019, Weisrock et al., 2018). This is especially important considering that, out of all vertebrate groups, amphibians are facing the most drastic population declines and extinction rates observed in the Anthropocene (Stuart et al., 2004, Hayes et al., 2010, Sparreboom, 2014, Lucas et al., 2024, Martel et al., 2013). Overall, by providing in the range of thousands to hundreds of thousands of high-quality SNPs, NewtCap facilitates the molecular study of salamandrids whilst whole genome sequencing of the gigantic genomes of salamanders remains unattainable.

## Acknowledgements

This project has received funding from the European Union’s Horizon 2020 research and innovation programme under the European Research Council grant agreement No. 802759 and the Marie Skłodowska-Curie grant agreement No. 655487. This work was performed using the compute resources from the Academic Leiden Interdisciplinary Cluster Environment (ALICE) provided by Leiden University. Daniele Canestrelli, Andrea Chiocchio, Michael Fahrbach, Willem Meilink, and Frank Pasmans contributed samples. The tissue sample of *Notophthalmus viridescens* (SMF 99040) was donated by the Senckenberg Research Institute and Nature Museum. The *Taricha* tissue samples were collected following protocols for animal welfare approved by UCLA and permitted by California Department of Fish and Wildlife (SCP # 12430). We would like to thank Erik-Jan Bosch for making the scientific drawings of the salamanders used in Figure 1 under the open content License of Naturalis Biodiversity Center (© CC BY-NC-ND 4.0), Benjamin Wiens for providing support with *triangulaR* adjustments, and Dr. Peter Scott for providing input on bioinformatic analyses. The *Triturus ivanbureschi* geographical map cut-out (Figure 4A) was made thanks to OpenStreetMap and its contributors (® CC BY-SA 2.0), and we obtained the *Taricha* cartoon from Figure 2 copyright free via user ‘clker-free-vector-images-3736’ through <u>Pixabay.com</u>.

## Data Accessibility and Benefit-Sharing Statement

The samples used in this study are in compliance with national laws and the Nagoya Protocol and detailed information can be found in Supplementary Materials, which are made publicly available on Zenodo (https://doi.org/10.5281/zenodo.15762260). Raw sequencing reads are accessible via BioProject ‘PRJNA1171613’ (https://www.ncbi.nlm.nih.gov/bioproject/PRJNA1171613). The main bioinformatic steps and scripts are provided on our GitHub (https://github.com/Wielstra-Lab/NewtCap_bioinformatics).

## Author Contributions

B.W., M.d.V., & J.F. conceived and designed the research. B.W., E.M., & H.B.S. initially designed the NewtCap probes for *Triturus*, and J.F. & B.W. further optimized the tool for broader use and higher efficiency. M.d.V., J.F., & A.T. performed the lab-work. M.d.V. conducted the pre-processing of the data, as well as all downstream analyses. G.B. contributed to the analyses of *Taricha*. M.d.V., B.W., & J.F. wrote the draft version of the manuscript, and all authors contributed to revising it.

## Supplementary Figures

**Figure S1:**
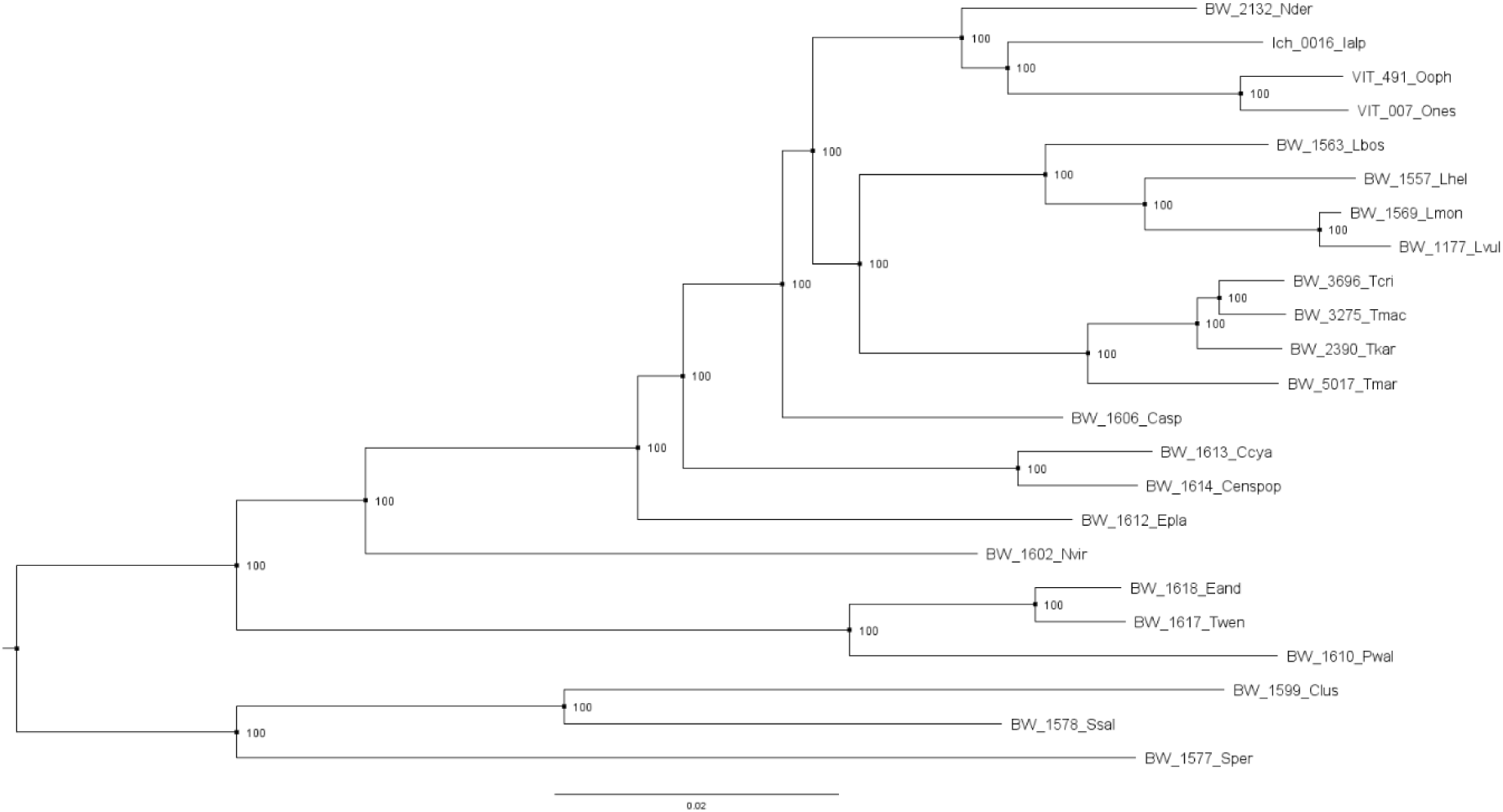
The raw, reconstructed NewtCap-based phylogeny of the Salamandridae family. *This is the same tree as provided in MS* Fig. 1, *but with original sample identifiers and bootstrap values. The tree is rooted on the branch separating the newts and the clade containing the true salamanders and* Salamandrina *(see also Fig. S2, which confirms the root position adopted here)*.

**Figure S2:**
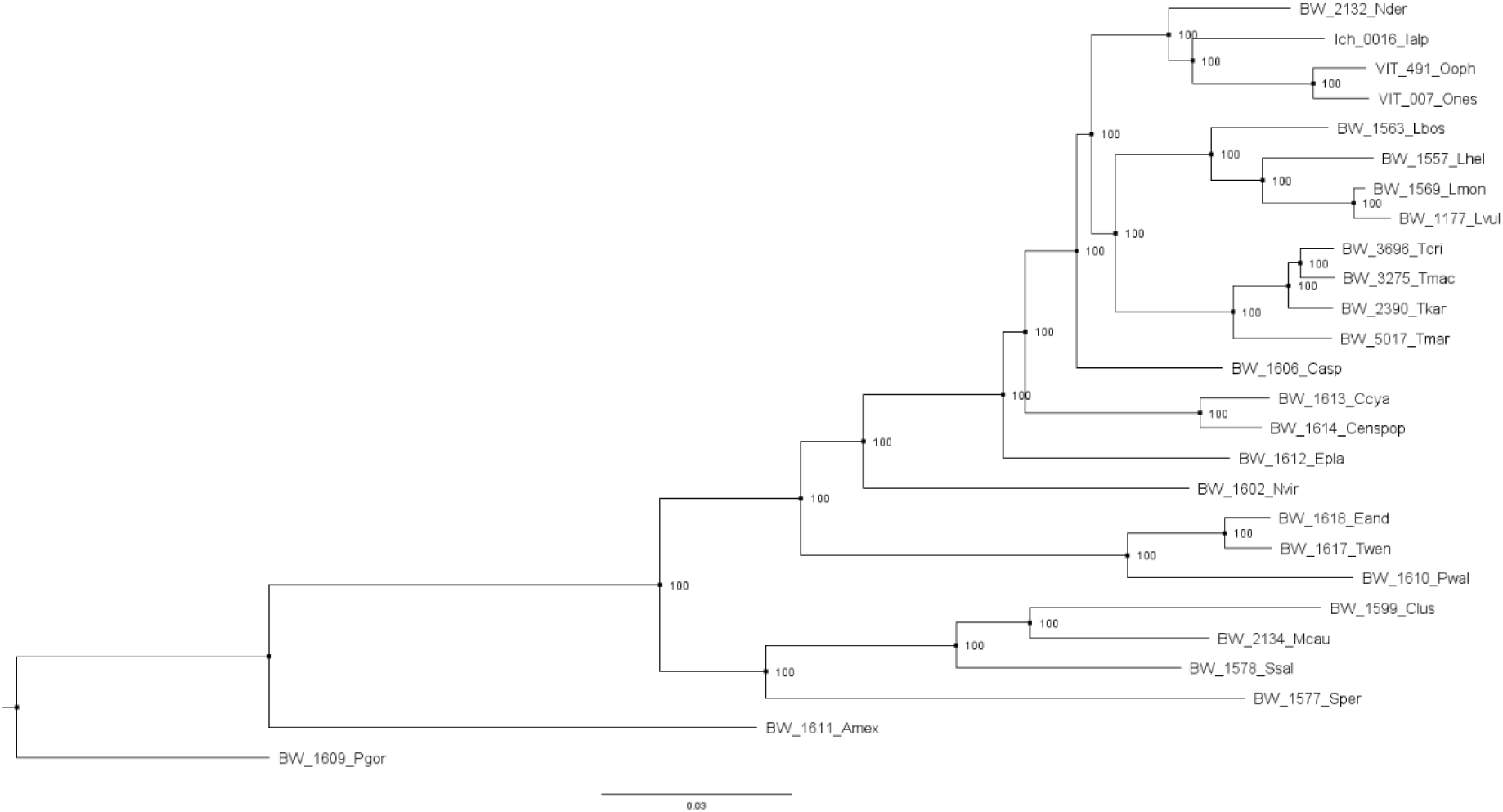
The raw, reconstructed NewtCap-based phylogeny of the Salamandridae family, including two non-salamandrids. *This tree is the result of an independent RAxML analysis that also includes a* Mertensiella caucasica *individual, and two non-salamandrid individuals: one* Ambystoma mexicanum *sample and one* Paradactylodon gorganensis *sample. The tree is based on 265,105 SNPs and shows sample identifiers and bootstrap values. The tree is rooted on the branch of* P. gorganensis, *which belongs to the Hynobiidae family and is more distantly related to Salamandridae than* A. mexicanum, *which belongs to the Ambystomatidae family (Marjanović and Laurin, 2013, Frost, 1985)*.

